# Clinical Benefit Index

**DOI:** 10.1101/473447

**Authors:** Amrita Basu, Laura Esserman

## Abstract

There are limited methods to analyze both efficacy of a drug and toxicity to a patient in response to treatment. We have developed a metric, the clinical benefit index (CBI) which is quantitative, and a scalable way to integrate patient-reported side effects and efficacy for adaptive randomization. Instead of using efficacy as a singular attribute in decision making for agent selection, we will consider the impact on QoL since many of the newer agents will be less toxic. This will be developed in effort to standardize PRO analysis methods. The clinical benefit index provides an alternative framework for evaluation and would help us to develop a regulatory model for approval of those drugs that have less toxicity by providing a replicable model for measuring toxicity and integrating it with efficacy. Reporting of clinical data can inform the management of toxicity and the integration into a clinical benefit index will help promote regulatory evaluation and preference for less toxic drug/combinations.

## 1 Clinical Benefit Index

We assume that have two measures, namely “Summary Score” and a “Response”. Response is based on clinical measurements while “Summary Score” is a metric based on patients’ answer to questions regarding their quality of life (QoL). The goal is to come up with a single clinical benefit index that captures both of these measures.

**Assumption:** We assume that for both of these measures, *the higher their values, the better it is for the patient*. Note that, if this weren’t true, one could always transform the measure(s) in question numerically to make it so.

### 1.1 Graphical Representation

Assume, without loss of generality, that the QoL metric and response are both on a scale of 0 to 1. This is for convenience only since any score could be standardized to ascertain the same. For that matter, we could also assume the distribution to be in the range of (*−∞, ∞*) and the subsequent duscussion will still go through. In the following, we represent the measures pictorially in Figure 1:

**Figure 1:**
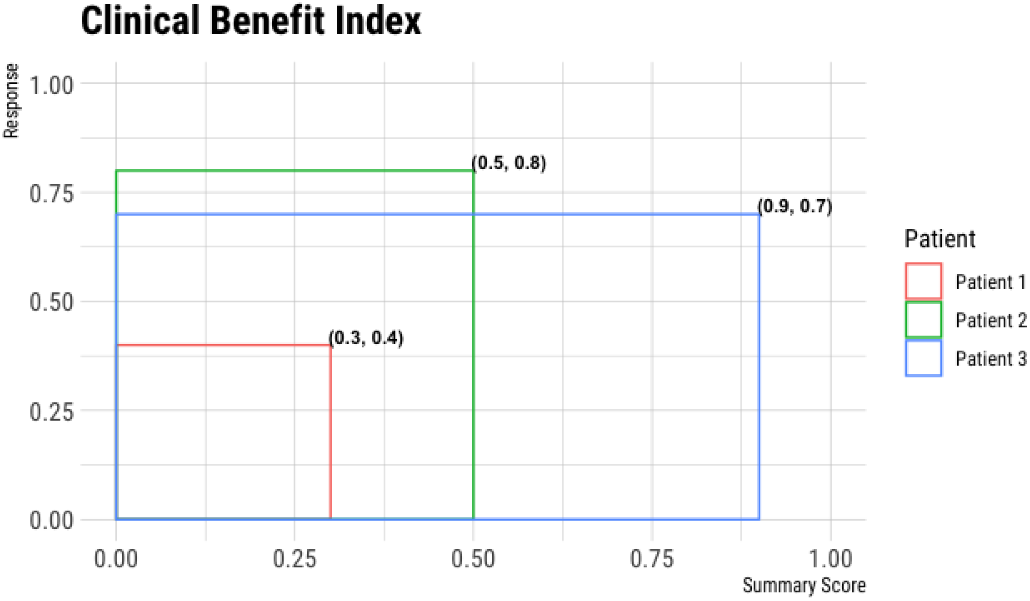
We show clinical benefit index calculation for three patients. The (Summary Score, Response) value-pair for the three patients are respectively, (0.3, 0.4), (0.5, 0.8) and (0.9, 0.7). The area of the red square is

#### 1.2

The two variables involved are `Response` and `Response` and both are assumed to have a domain of distribution in [0, 1]. These two variables are distributed jointly on a [0, 1] × [0, 1] rectangular surface. We assume that the joint distribution is a bivariate uniform over the rectangale. Mathematically, the bivariate probability density function (pdf) could be written as:

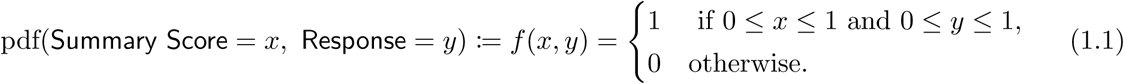

For each patient, and for each survey, our proposal is to calculate the area that the the patient covers in the [0, 1] × [0, 1] rectangale. As an example, assume that patient P has the value

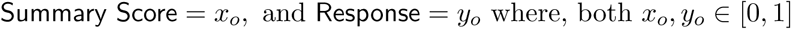

The area covered by the point (*x_0_, y_0_*) in the rectangle has the interpretation that,

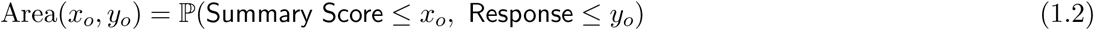

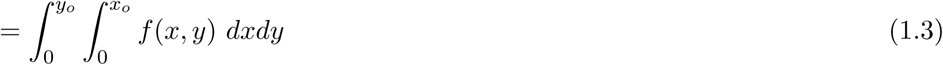

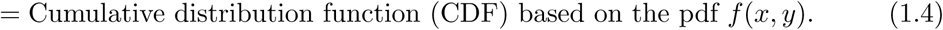

#### Remark 1.1

The above calculation is based on the assumption that the bivariate distribution is uniform. In reality, this distribution could be bivariate Gaussian, or some other heavy-tailed distribution; we can accomodate all these different cases as well by approximating the density empirically from past data.

